# A 23-copy tandem repeat of a cytochrome P450 gene underlies an adaptive introgression breakthrough in *Ciona intestinalis*

**DOI:** 10.64898/2026.10.01.755871

**Authors:** Manon Le Goff, Fabienne Justy, Theo Deremarque, Frédérique Viard, Nicolas Bierne

## Abstract

Introgression between highly divergent species is generally selected against, yet some foreign alleles can break through species barriers when they provide a sufficient selective advantage, a process known as adaptive introgression. Here, we characterize the structural variant underlying adaptive introgression from *Ciona robusta* into *C. intestinalis*, two species separated by 14.5% net molecular divergence. Previous work identified a narrow peak of *C. robusta* ancestry on chromosome 5 that reached high frequency in several *C. intestinalis* populations despite otherwise restricted introgression. Using whole-genome Nanopore sequences from 15 *C. intestinalis* and three *C. robusta* individuals, together with short-read datasets including parent-offspring trios, we narrowed down the core introgressed segment from 16 to three genes and characterized copy-number variation affecting the cytochrome P450 gene CYP2U1c. The introgressed haplotype carries a tandem amplification of a 5,173-bp segment containing CYP2U1c, with 23 consecutive copies and conserved repeat boundaries. The nearly 120-kb array could not be fully resolved by long-read sequencing, but coverage consistently supported approximately 23 copies per introgressed haplotype. Despite its extreme size and repetitive structure, the allele showed Mendelian inheritance. CYP2U1c copy number was also polymorphic in *C. robusta*. Long reads fully resolved two structurally distinct haplotypes in one individual, carrying either a duplication encompassing adjacent CYP2U1b and CYP2U1c genes or five CYP2U1c copies. Their distinct breakpoints indicate independent origins. Our results reveal independently arising CYP2U1c amplifications in the donor species and show that an extreme high-copy-number allele was subsequently favoured following introgression into *C. intestinalis*.

## INTRODUCTION

Introgression is now recognised as an important source of genetic variation in many species (Schumer & Rieseberg, 2026). Although gene flow between divergent lineages is often selected against, introgressed alleles can occasionally confer a selective advantage in the recipient population, a process referred to as adaptive introgression (Hedrick, 2013). Most studies of adaptive introgression have focused on single-nucleotide polymorphisms (SNPs), although the contribution of structural variants (SVs) is now receiving increasing attention (Yan *et al*., 2021; Hsieh *et al*., 2026). This partly reflects recent technological advances, as SVs have historically been difficult to detect and characterize using short-read sequencing, especially in repetitive regions (Ho *et al*., 2020). Long-read sequencing technologies now allow more direct characterization of SVs, including large and complex variants that are difficult to resolve with short reads (Amarasinghe *et al*., 2020; Edwards & Li, 2026).

SVs are increasingly recognized as an important source of genetic and phenotypic variation, with potentially important roles in adaptation (Wellenreuther *et al*., 2019; Mérot *et al*., 2020). Among SVs, copy number variations (CNVs) can have particularly strong phenotypic effects by altering gene dosage and expression, or by generating redundant gene copies that may subsequently diverge in function (Freeman *et al*., 2006; Zhang *et al*., 2009; Ohno, 2013). CNVs can arise through several distinct mechanisms. First, replication errors or DNA repair can duplicate a genomic segment, for example through template switching during replication, without requiring recombination between homologous chromosomes (Hastings *et al*., 2009). Second, non-allelic homologous recombination, also called ectopic recombination, can occur when similar repeated sequences located at different genomic positions misalign during meiosis. An unequal crossover between homologous chromosomes then produces reciprocal products, with a duplication of the intervening segment on one chromosome and a deletion on the other (Hastings *et al*., 2009). Once a duplication is present in tandem, the duplicated copies themselves provide homologous sequences that can misalign during subsequent recombination events. Unequal crossing-over can then increase the number of copies on one chromosome while decreasing it on the other, allowing tandem arrays to rapidly expand or contract over generations (Elemento *et al*., 2002).

Such changes in copy number have repeatedly been involved in adaptation, including responses to environmental changes and anthropogenic selection (Kondrashov, 2012). Gene amplification, for example, is an important mechanism of insecticide resistance in mosquitoes, where increased copy number can increase the expression of resistance genes (Raymond *et al*., 2001; Nagi *et al*., 2024). CNVs may therefore constitute large-effect adaptive variants that can also be transferred between divergent lineages through introgression. In *Uria* seabirds, introgressed CNVs have been associated with adaptation to cold environments (Tigano *et al*., 2018). More recently, Villanea *et al*., (2025) showed that a positively selected archaic MUC19 haplotype in humans carries a higher copy number of a variable number tandem repeat, linking introgression, tandem copy-number variation and positive selection. These examples show that adaptive introgression can involve not only nucleotide variants, but also large and structurally complex alleles whose characterization has only recently become accessible.

In this context, we are interested in anthropogenic introgression, which results from introgression caused by human-induced contact between divergent lineages (McFarlane & Pemberton, 2019; Viard *et al*., 2020). Fraïsse *et al*., (2022) provided a striking example in the two sea squirt species *Ciona robusta* and *C. intestinalis*. Despite their high divergence, with 14.5% net molecular divergence, Fraisse et al. (2022) identified introgression from the introduced non-indigenous species *C. robusta* into the *C. intestinalis* genome, restricted to a single genomic region on chromosome 5. Within this narrow breakthrough of an otherwise strong barrier to introgression, long introgressed tracts reached high frequencies in several populations, providing evidence for adaptive introgression (Fraïsse et al. 2022; Le Goff et al. 2026). The stability of the introgressed allele frequency over a 20-generation interval (Touchard et al. 2024), together with evidence from the joint distribution of introgressed tract lengths and allele frequencies (Le Goff et al. 2026), further suggests that the introgressed allele is under local selection. Fraisse et al. (2022) localized the peak of introgression to a core region containing 16 genes. Among these, a cytochrome P450 gene annotated as CYP2U1 emerged as a candidate because SNPs within this gene showed unusually high coverage and unbalanced allelic coverage ratios, suggesting the presence of multiple gene copies. However, the precise target of selection and the nature of this putative copy number variant remained unresolved.

Here, we aimed to refine the target of selection within the introgressed region and to characterize the putative structural variant associated with CYP2U1. In particular, we tested whether the adaptive *C. robusta* allele corresponds to a large tandem amplification of this gene and whether this complex allele is stably inherited. To address these questions, we combined newly generated and previously published genomic data. New data consisted of Nanopore long-read sequencing for 15 *C. intestinalis* and 3 *C. robusta* from 5 localities. We complemented these data by reanalysing previously generated genomic datasets, including Nanopore reads from two individuals used to produce chromosome-scale genome assemblies (Satou et al., 2021), and short-read data from two phased-genome studies, one based on parent-offspring trios (Fraisse et al., 2022) and the other on a linked-read sequencing technique, namely haplotagging (Le Goff et al., 2026). Together, these complementary datasets allowed us to refine the introgressed region, characterize the structure and copy number of the CYP2U1 amplification, and examine its inheritance. We show that the peak of adaptive introgression can be narrowed down to three genes only in the reference genome, including two genes annotated as CYP2U1, and that the introgressed allele carries a tandem array of 23 highly similar copies, with an uncertainty of one copy, of one of these two CYP2U1 genes. Despite its large and repetitive structure, the tandem array allele follows Mendelian inheritance. These results identify a large, likely recent, gene amplification as the target of adaptive introgression between two highly divergent species.

## MATERIALS AND METHODS

### Long-read sequencing and data processing

#### Sampling, DNA extraction and Nanopore sequencing

To characterize the genomic structure of the adaptive introgression region, we generated long-read sequencing data from 15 *C. intestinalis* and 3 *C. robusta* individuals. Individuals selected for this study were sampled during late summer 2021 from three localities: Brixham (Northern English Channel, UK), Brest-Moulin Blanc (Southern English Channel, France) and Arcachon (Bay of Biscay, France). Individuals had previously been analysed using a KASP genotyping assay that includes a diagnostic locus located at the core of the introgression region on chromosome 5 (Touchard et al. 2024). At all three locations, the frequency of introgression was high (>65%), with the three genotypic classes represented: II *C. intestinalis* homozygotes, IR heterozygotes, and RR homozygotes for the introgressed *C. robusta* allele (Touchard et al. 2024). Approximately 10 individuals previously analysed by Touchard et al. (2024) were selected from each locality. Tissues preserved in RNAlater were used to extract genomic DNA using the Monarch HMW DNA Extraction Kit for Tissue (New England BioLabs). DNA quality and concentration were assessed using a NanoDrop 2000 spectrophotometer (Thermo Fisher Scientific, USA) and a Qubit 3.0 fluorometer (Life Technologies, USA). For DNA showing degradation, the SRE XS kit (Circulomics, US) was used to improve quality before library preparation. Fifteen *C. intestinalis* individuals and three *C. robusta* individuals with the highest DNA quality and quantity were chosen for ONT long-read sequencing. Libraries were prepared using the ONT Template Prep Kits SQK-LSK110 and SQK-LSK112 according to standard protocols and sequenced on the ONT MinION platform using R9.4.1 flow cells (Oxford Nanopore Technologies, Oxford, UK). In addition to the long-read data generated in this study, we used Nanopore data previously generated by Satou et al. (2021) for two *C. intestinalis* individuals, one from Plymouth (English Channel, UK; GCA_018327805.1) and one from Roscoff (English Channel, France; GCA_018327825.1).

#### Sequence processing and mapping

Initial basecalling of the newly generated Nanopore data was performed using ONT’s Dorado basecaller (https://github.com/nanoporetech/dorado). Input data were converted from pod5 to fast5 format. The resulting sequences were mapped to the *C. robusta* reference genome (GCA_009617815.1, Satou et al. 2019). This genome is referenced under the name *C. intestinalis* type A, the name used for *C. robusta* before the taxonomic status of the two taxa was re-evaluated (Brunetti et al. 2015). Mapping was performed using Dorado, which uses the minimap2 aligner (Heng 2018). The published Plymouth and Roscoff Nanopore reads were similarly mapped to the *C. robusta* reference genome using minimap2. Alignments were sorted and indexed using samtools (Danecek et al. 2021), and data quality was assessed using samtools and overall statistics generated with MultiQC (Ewels et al. 2016). Sequencing depth was retrieved from the alignments using samtools.

### Data analyses

#### Delineation of the core adaptive introgression interval

In order to better delineate the introgression region, and thereby narrow down the adaptive introgression interval identified by Fraisse et al. (2022), we reanalysed local ancestry estimates previously obtained by Le Goff et al. (2026). Fraisse et al. (2022) used haplotypes phased by descent to identify the core of the introgression peak, which encompassed 16 genes. The larger sample available from Le Goff et al. (2026) should capture additional recombination breakpoints and allow this interval to be narrowed down further. The Le Goff et al. (2026) dataset comprised 96 individuals from seven localities, physically phased using haplotagging, a linked-read sequencing method that relies on tagging long DNA molecules prior to sequencing (Meier et al. 2021). Local ancestry had been reconstructed for each haplotype using LOTER (Dias-Alves et al. 2018) by Le Goff et al. (2026). Here, we used these local ancestry estimates to identify the interval along chromosome 5 where the proportion of *C. robusta* ancestry reached its maximum, which we considered as the core adaptive introgression interval expected to contain the target of selection.

#### Copy-number estimation from sequencing coverage

The copy number of the segment containing the candidate CYP2U1 gene was assessed using both long- and short-read sequencing data. Hereafter, CYP2U1 refers specifically to the candidate gene located within this segment, rather than to the other CYP2U1 genes present in the genome. Long-read data included the Nanopore data generated in this study and the two published *C. intestinalis* Nanopore datasets from Satou et al. (2021). Short-read data included the 96 individuals from Le Goff et al. (2026) and the parent-offspring dataset produced by Fraisse et al. (2022). For each individual, the average coverage of the CYP2U1-containing segment (positions 1,049,202–1,054,374 bp in the *C. robusta* reference genome) was calculated and divided by the expected diploid coverage. Because the denominator represents coverage from both parental haplotypes, the resulting ratio was multiplied by two to obtain the total number of copies across the two haplotypes. We tested several approaches to estimate the expected diploid coverage used as the denominator to calculate relative sequencing depth. We compared mean coverage across all chromosomes, mean coverage within the introgression region excluding the CYP2U1-containing segment (positions from 1 Mb to 1,049,202 bp and from 1,054,374 bp to 1.1 Mb), and mean coverage across genic regions identified using BUSCO (v6.1.0, Tegenfeldt *et al*., 2025). We also assessed the influence of the reference genome by comparing relative coverage estimates obtained after alignment to the *C. robusta* and *C. intestinalis* reference genomes.

#### Inheritance of the CYP2U1 copy-number variant

To examine the inheritance of the CYP2U1 copy-number variant, we reanalysed short-read sequencing data from the parent-offspring crosses of Fraisse et al. (2022). In that study, laboratory crosses were performed using individuals from the Aber Wrac’h and Moulin Blanc marinas (English Channel, France), resulting in six interspecific and ten intraspecific crosses, including six between *C. intestinalis* individuals and four between *C. robusta* individuals. Parents and offspring were sequenced using 100-bp paired-end Illumina reads, and genotypes at the introgression region were phased by transmission. Here, we used the available FASTQ data, mapped them to the *C. robusta* reference genome (GCA_009617815.1, Satou et al. 2019) using BWA-MEM (Li, 2013), and estimated CYP2U1 copy number from coverage as described above. Individuals with insufficient sequencing coverage were excluded, resulting in some incomplete crosses. Copy-number estimates were then compared between parents and offspring to examine the inheritance of the CYP2U1 copy-number variant.

#### Structural characterization of the CYP2U1 tandem repeat

To study the structure and relationships among copies of the candidate CYP2U1 gene, we used our Nanopore data. The segment corresponding to the candidate CYP2U1 is present as a single copy in the *C. robusta* reference genome. This sequence was mapped to the two *C. intestinalis* genome assemblies to examine the structure and copy number of the CYP2U1 locus. We also retrieved long reads spanning the region of interest for each *C. intestinalis* individual. We then focused on the Roscoff assembly, in which 21 tandem copies of the candidate CYP2U1 were identified. The sequence containing the tandem repeat was extracted from the assembly and compared with long reads from the same individual to characterize the structure and boundaries of the repeat and assess whether the assembled repeat structure was supported by the raw reads. The candidate CYP2U1 gene and its flanking sequence were mapped to both the assembled sequence and the spanning reads. The 21 CYP2U1 copies identified in the Roscoff assembly were extracted and aligned using MAFFT (Katoh *et al*., 2019), and a Neighbor-Joining tree was constructed from a distance matrix based on pairwise nucleotide differences. For building the tree, we used two external groups: the CYP2U1 sequence from the *C. robusta* reference genome and a *C. intestinalis* consensus sequence generated by calling variants from an II individual against the *C. robusta* reference genome. We also examined one of the newly sequenced *C. robusta* individuals for which coverage analysis indicated multiple copies of the CYP2U1-containing segment. Long reads spanning the region of interest were retrieved for this individual, and their structure was characterized by mapping the candidate CYP2U1 sequence from the *C. robusta* reference genome.

## RESULTS

### The core adaptive introgression interval is narrowed down to three candidate genes, including an amplified CYP2U1 gene

We used the ancestry profiles obtained from the haplotagging dataset to identify the interval with the highest proportion of *C. robusta* ancestry within the *C. intestinalis* genomes. This narrowed down the core introgression interval to a small segment between positions 1,042,729 bp and 1,055,324 bp of the *C. robusta* reference genome (Fig. 1B), containing only three genes.

**FIGURE 1.**
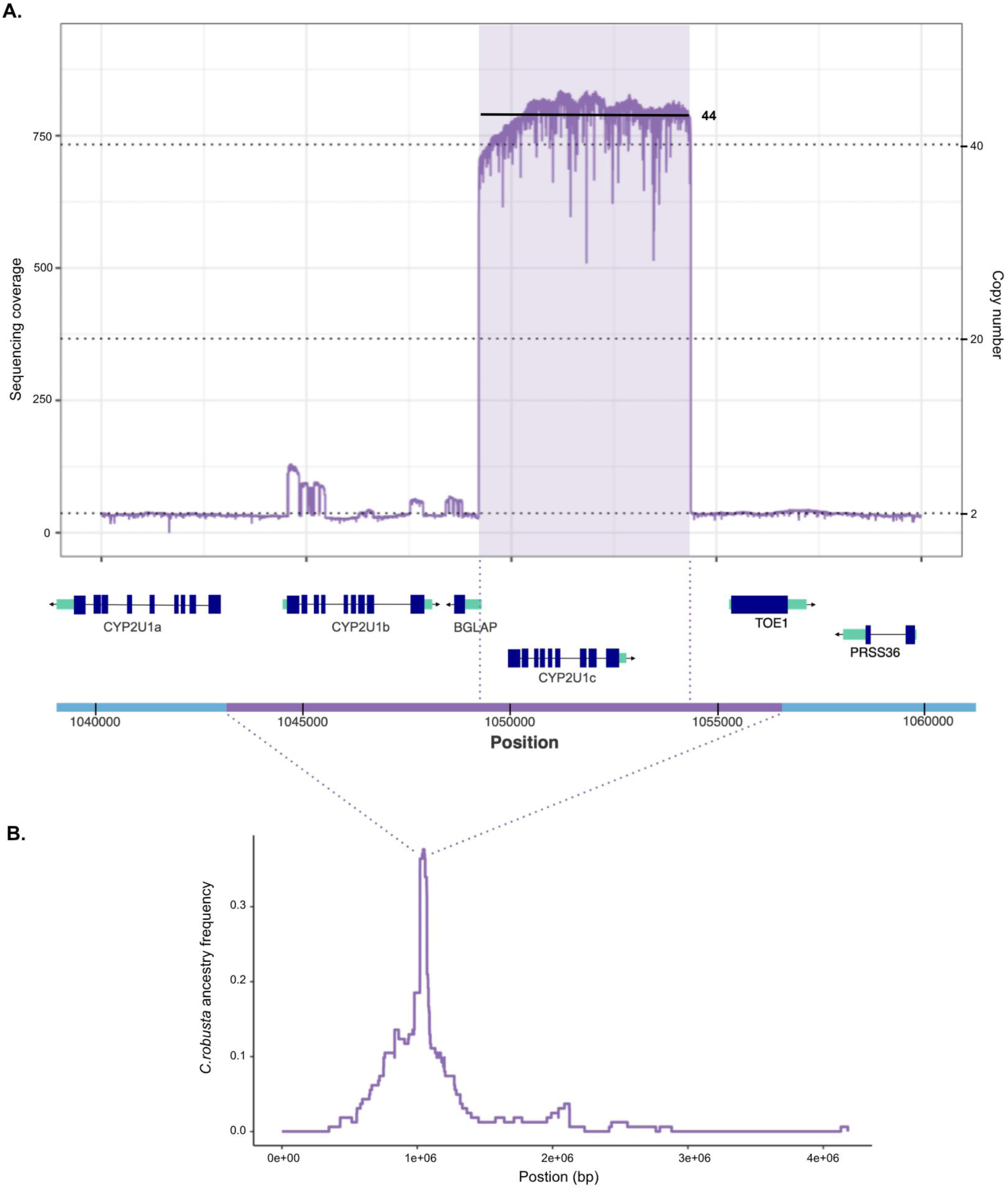
Focus on the region under selection. A) Example of sequencing coverage along the genome for one individual. Genes within the region are shown below, together with the interval of maximum *C. robusta introgression*. Coverage profiles for all other individuals are shown in Supplementary Fig. S1. B) Frequency of C. robusta ancestry along the genome across all individuals, inferred from phased short-read data (Le Goff et al. 2026).

Annotation revealed two cytochrome P450 family 2 subfamily U member 1 (CYP2U1) genes and one bone gamma-carboxyglutamate protein (BGLAP) (Fig. 1A). A third CYP2U1 gene is located immediately upstream. We hereafter refer to the three CYP2U1 genes as CYP2U1a, CYP2U1b and CYP2U1c, according to their order along the chromosome (Fig. 1A). CYP2U1c is located within a 5174 bp segment (positions 1,049,201bp to 1,054,375bp) showing substantially higher sequencing coverage (Fig. 1A), suggesting that the entire segment is present in multiple copies. The sharp drop in coverage at both ends of this segment further suggests that the repeated copies have conserved boundaries, with consistent breakpoint positions across individuals (Supplementary Fig. S1).

### Long-read coverage provides more reliable estimates of CYP2U1c copy number

Copy-number estimates from short-read data showed substantial biases. First, for the short-read dataset of Le Goff et al. (2026), non-introgressed II homozygotes yielded an average estimate of approximately six copies (Fig. 2A), whereas two copies were expected, one on each chromosome. Conversely, copy number was underestimated in introgressed RR homozygotes. These individuals had an average estimated copy number of approximately 40, rather than twice the approximately 23 copies estimated in IR heterozygotes (Fig. 2A). We therefore did not consider absolute copy-number estimates from this dataset to be reliable.

**FIGURE 2.**
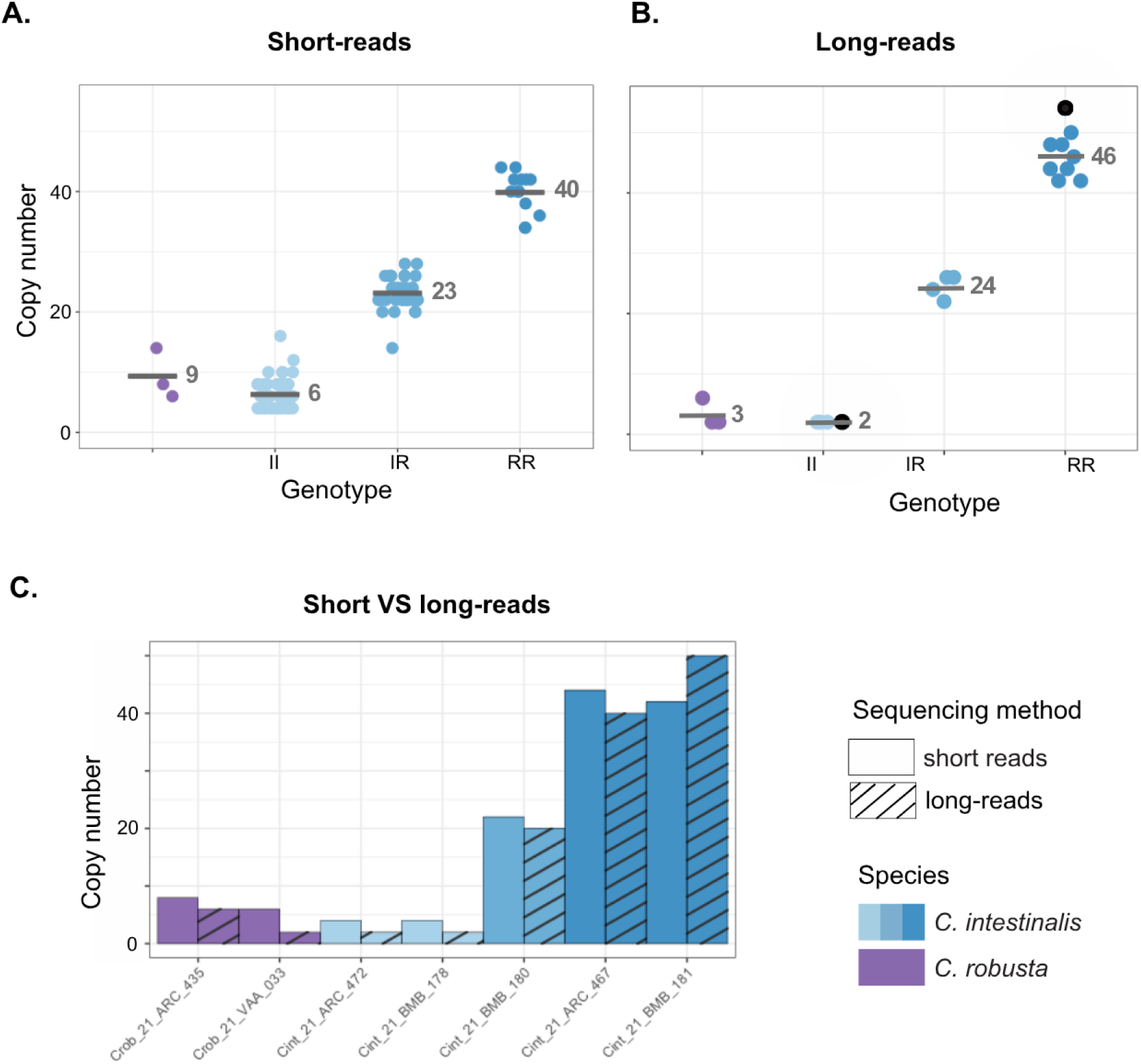
Number and distribution of CYP2U1 gene copies in *Ciona robusta* and *C. intestinalis*. Copy number was estimated by normalizing CYP2U1 coverage to the average coverage across all chromosomes except chromosome 5. A) Copy-number estimates for all individuals from the short-read dataset (Le Goff et al. 2026). B) Copy-number estimates for all individuals from the long-read dataset. Black dots indicate the two assembled genomes from Satou et al. (2021). C) Comparison of copy-number estimates obtained from short- and long-read data.

Nevertheless, despite being a short-read sequencing dataset, coverage of the repeated segment in the parent-offspring dataset of Fraisse et al. (2022) clearly correlated with the previously determined II, IR and RR genotypes, which had low, intermediate and high coverage, respectively (Fig. 3). However, copy-number estimates differed between individuals heterozygous for the copy-number variant according to their species background. *C. robusta* heterozygotes consistently showed lower copy-number estimates than *C. intestinalis* heterozygotes. Copy-number estimates for RR homozygotes were also inconsistent with those of IR individuals, as RR homozygotes should carry approximately twice as many copies of the repeated allele as IR individuals. Despite this variability, we observed a perfect correspondence between the genotypes of the parents and those of the offspring, supporting a Mendelian inheritance of the copy-number variant (Fig. 4B). This held true for both intraspecific and interspecific crosses. These results suggest that biases in short-read mapping depend on the genomic background, possibly because reads from the repeated segment can map ambiguously among the numerous CYP-related sequences present in the genome.

**FIGURE 3.**
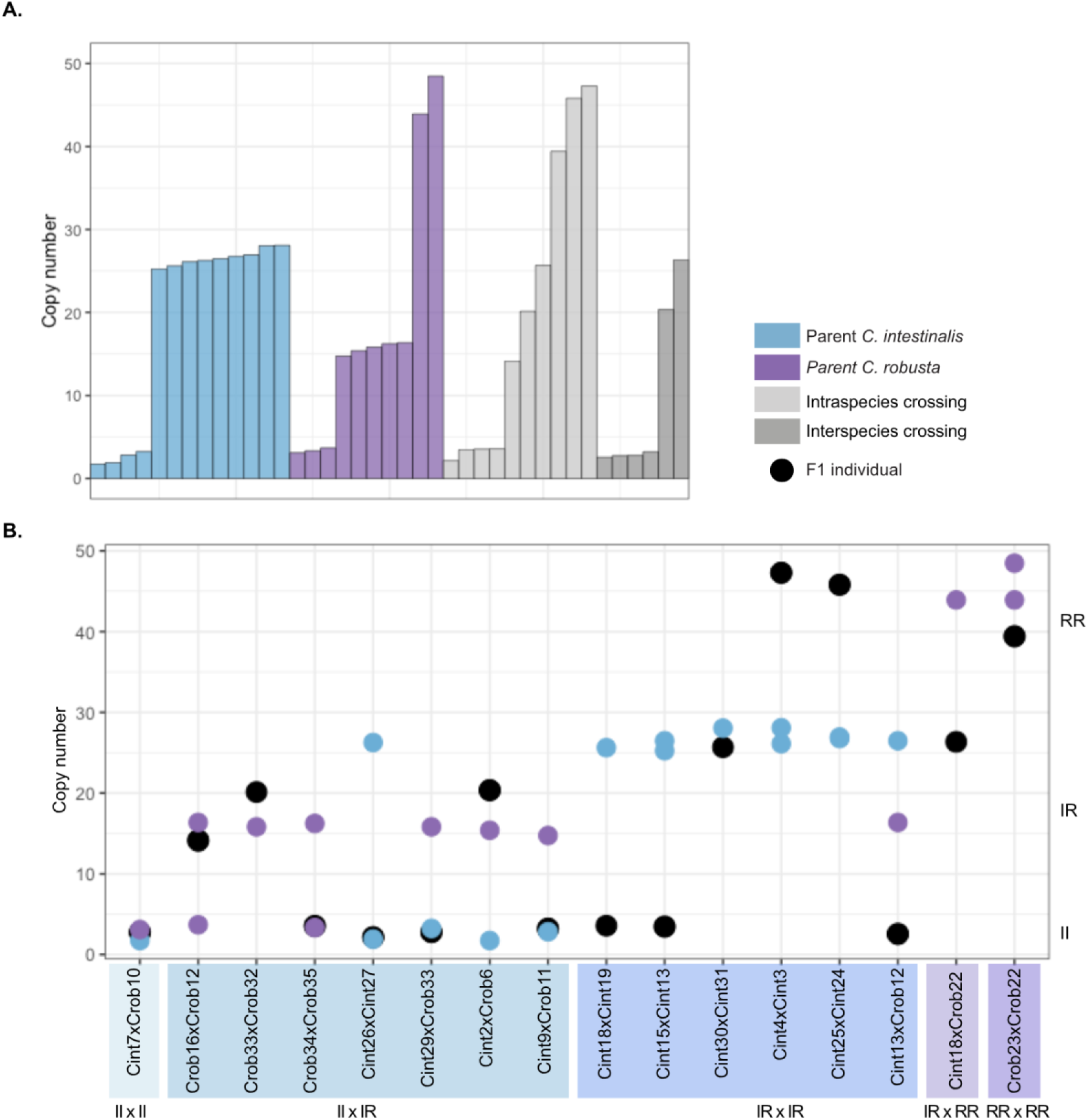
Number of copies in the parent-offspring crosses dataset. A) Copy-number estimates for all individuals according to species and cross type. B) Copy-number estimates for each offspring and its parents. Point colors indicate parental species. Colors of the crosses represent their type: II × II, II × IR, IR × IR, IR × RR, and RR × RR, where II and RR are homozygotes and IR are heterozygotes.

**FIGURE 4.**
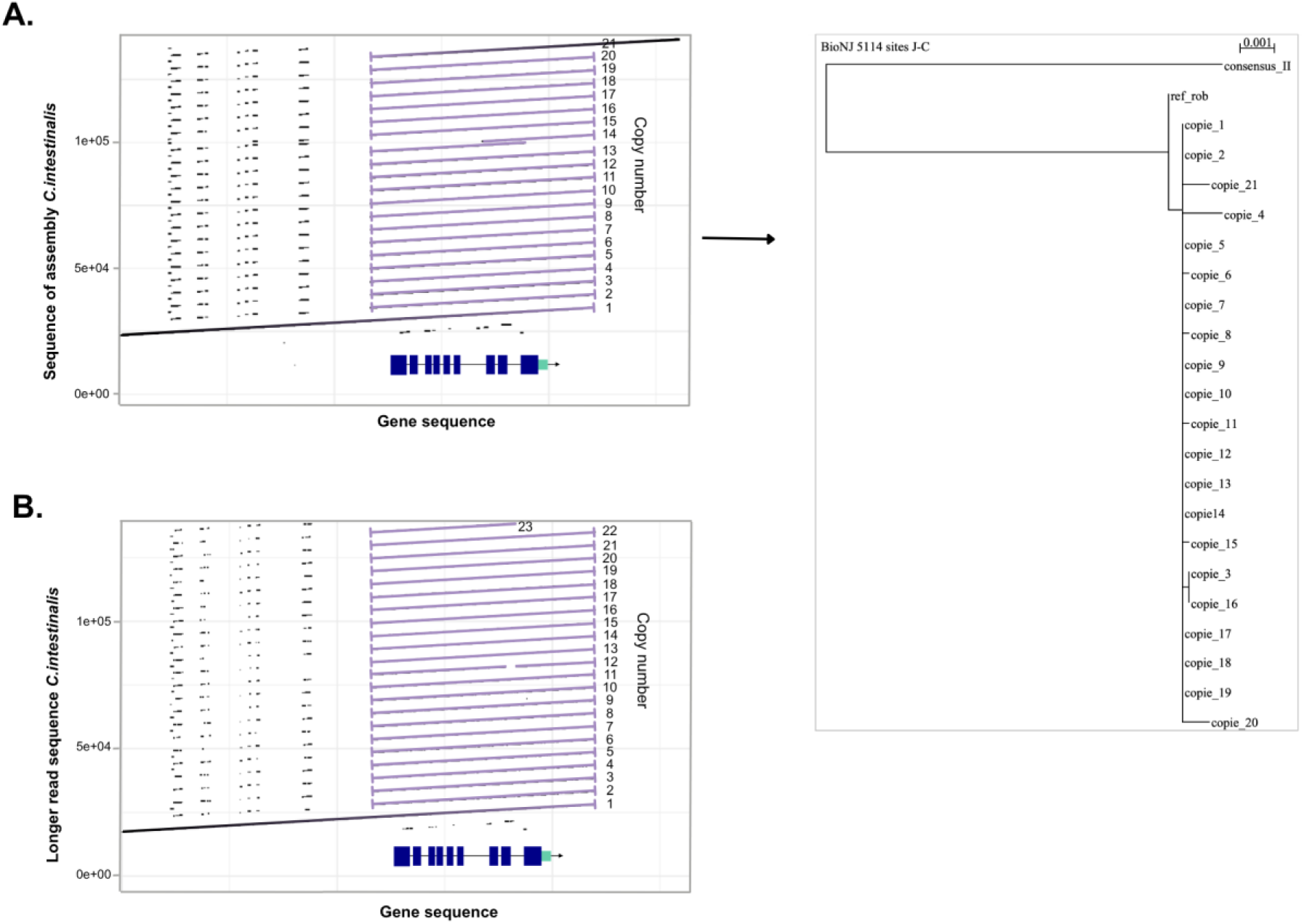
Structural variation in the CYP2U1 gene. A) Dot plot of the Roscoff *C. intestinalis* reference assembly (Satou et al. 2021), restricted to the region of interest and aligned against the CYP2U1c gene region. The 21 copies of the CYP2U1c containing segment identified in the assembly are shown below the dot plot across the two scaffolds used to assemble this region. The tree on the right was obtained from the alignment of these 21 copies, together with the *C. robusta* reference copy and the consensus copy from *C. intestinalis* individuals lacking the introgressed allele (II genotype) as outgroups. B) Dot plot of the longest read containing 23 consecutive copies identified in the Roscoff *C. intestinalis* long-read dataset, aligned against the CYP2U1c gene region.

Long-read data provided much more consistent copy-number estimates, likely because long reads can be mapped more accurately to their true genomic position. Across our different denominators, the expected ratio between II, IR and RR genotypes was recovered (Fig. 2B, Supplementary Fig. S2).

II homozygotes consistently had an estimated copy number of two across individuals and denominators. IR heterozygotes were found to carry half the number of repeated copies found in RR homozygotes, plus one copy that corresponds to the single CYP2U1 copy carried by the non-introgressed haplotype. This result was recovered except when coverage within the introgressed region itself was used as the denominator (Supplementary Fig. S2), in which case IR heterozygotes were underestimated by approximately one copy. However, none of the three denominators substantially reduced the variability in estimated copy number among IR and RR individuals. Copy numbers also varied among *C. robusta* individuals. In contrast to the *C. robusta* individuals of the study of Fraisse et al. (2022), none of the three long-read sequenced *C. robusta* individuals carried the 23-copy allele, either in the heterozygous or homozygous state. Two individuals were homozygous for a one-copy allele, whereas the third had an estimated six copies overall (individual Crob_21_ARC_435 in Fig. 2C). The distribution of these copies between the two haplotypes is investigated further below.

### Structural characterization of the introgressed CYP2U1c tandem repeat and of a distinct structural variant in one *C. robusta* individual

We next investigated the structure of the CYP2U1c copy-number variation using long-read data and genome assemblies. Coverage analysis of the two *C. intestinalis* individuals sequenced by Satou et al. (2021) identified the Plymouth individual as II and the Roscoff individual as RR. We therefore focused on the Roscoff genome assembly, for which approximately 23 copies of CYP2U1c were expected on each parental chromosome based on the coverage analysis. Mapping the CYP2U1c sequence from the *C. robusta* reference genome to the Roscoff assembly identified 21 copies arranged in tandem. All copies shared the same repeat boundaries, confirming that the amplified segment was assembled as a tandem repeat. The repeated segment extended from positions 1,204,193 to 1,313,027 bp on chromosome 5 of the Roscoff *C. intestinalis* assembly (Fig. 4A). CYP2U1c exons also showed sequence similarity to the upstream CYP2U1b gene, but this similarity did not extend to the introns. The 21 assembled CYP2U1c copies showed very little sequence variation, with pairwise sequence similarities ranging from 99.99% to 100% (Fig. 4A). In the phylogenetic tree, the *C. intestinalis* consensus sequence showed substantially more nucleotide differences from the repeated copies, whereas the *C. robusta* reference CYP2U1c sequence was more similar to them.

However, the 21-copy structure of the assembly could not be validated directly from the raw Nanopore reads. None of the reads from the Roscoff individual spanned the tandem repeat from one end to the other (Supplementary Fig. S3). The longest read overlapping the repeat was 138,367 bp long and extended to the 23rd CYP2U1c copy without reaching the opposite end of the repeat (Fig. 4B). Other reads contained smaller numbers of consecutive CYP2U1c copies, but no single read supported a complete 21-copy allele. Sequence variation among copies could not be reliably estimated from these reads because of the relatively high error rate of the Nanopore chemistry used, whereas sequence errors were largely corrected during genome assembly. Consequently, although the raw reads confirm the presence of a long CYP2U1c tandem repeat, they do not allow us to determine whether the 21-copy structure in the assembly represents one of the two haplotypes. In particular, the coverage estimate could be explained by two alleles carrying approximately 23 copies each, or by different copy numbers on the two chromosomes, for example a 21-copy allele together with an allele carrying approximately 25 copies.

We then examined the single *C. robusta* individual showing increased CYP2U1c copy number in our Nanopore dataset (Crob_21_ARC_435 in Fig. 2C). The standard coverage-based method based on the boundaries of the introgressed CYP2U1c tandem repeat gave an estimated total copy number of 5.6, which we rounded to six copies.

However, coverage along the chromosomal region showed a series of distinct plateaus of coverage rather than the sharp boundaries observed for the introgressed tandem repeat in *C. intestinalis* (Fig. 5A), indicating a more complex structural variant. The highest coverage corresponded to approximately seven copies, with additional plateaus corresponding to approximately two, three and five copies. Because this *C. robusta* variant contained fewer repeats, some individual long reads spanned the entire structural variant and allowed its organization to be reconstructed (Supplementary Fig. S4). Two types of spanning reads were recovered, corresponding to the two parental haplotypes (Fig. 5B, Supplementary Fig. S4). One haplotype carried a duplication of a larger segment encompassing CYP2U1b, BGLAP and CYP2U1c. It therefore contained two copies of CYP2U1b, two copies of BGLAP (gene annotated between the two CYP2U1) and two copies of CYP2U1c. The second haplotype carried five copies of a segment containing BGLAP and CYP2U1c. Together, these two haplotypes account for the successive coverage plateaus observed across the region, with two copies of a segment containing CYP2U1b, BGLAP and CYP2U1c on one chromosome and five copies of a segment containing BGLAP and CYP2U1c on the other, the duplicated segments having different boundaries in the two haplotypes (Fig. 5B).

**FIGURE 5.**
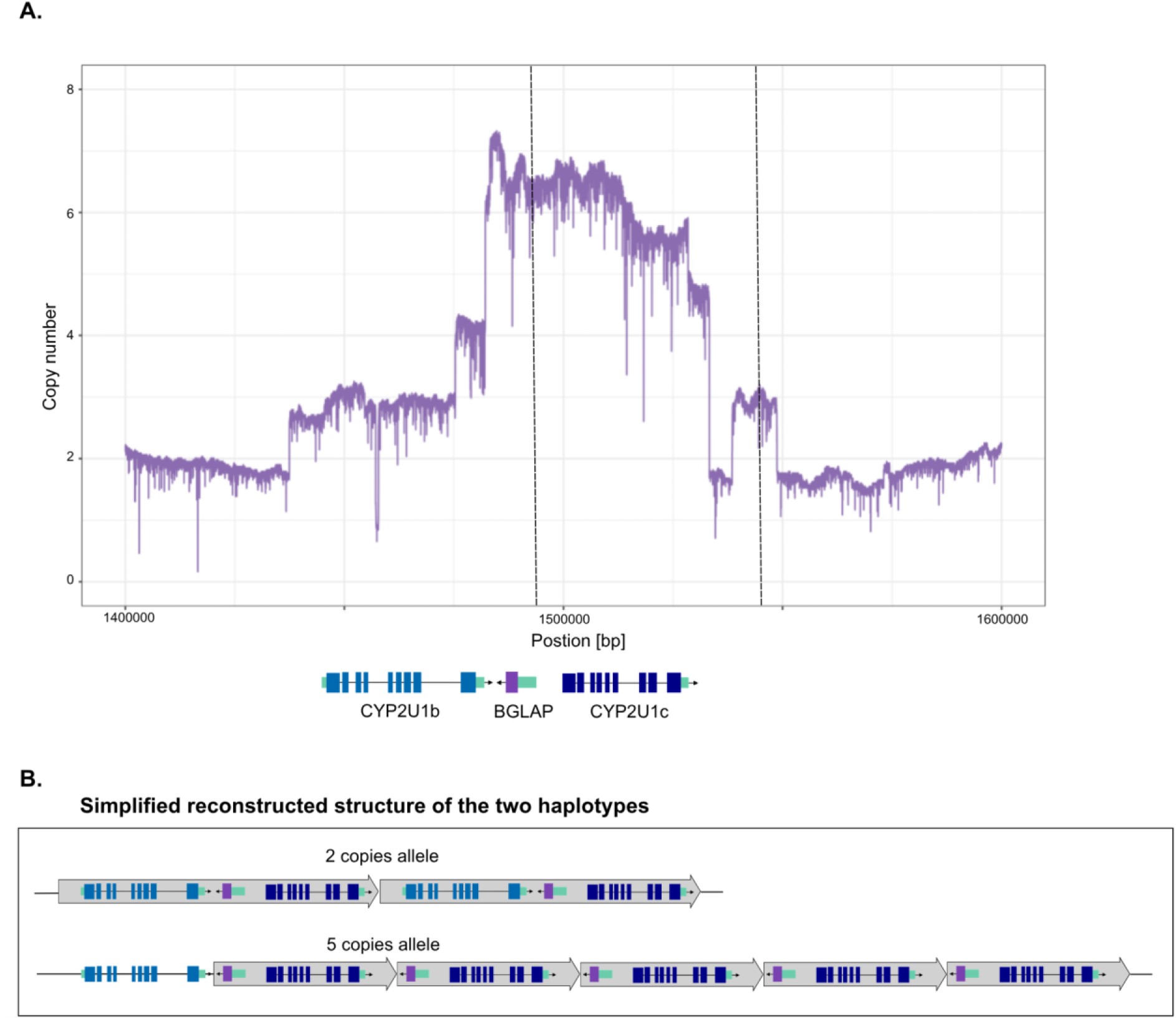
Copy-number variation in a C. robusta individual. A) Copy-number estimates based on long-read sequencing coverage for the *C. robusta* individual with multiple CYP2U1 copies. Dashed lines delimit the repeated segment identified in *C. intestinalis*. B) Schematic representation of the two haplotypes inferred from individual long reads spanning the entire structural variants (Supplementary Fig. S4).

During the exploration of individual long reads we noted that some reads showed additional structural configurations, including inverted rather than tandem duplications (Supplementary Fig. S5), which initially made the structure of the locus difficult to resolve. However, these configurations were not consistent across reads and were incompatible with the two parental haplotypes reconstructed above, which together accounted for the observed coverage profile. We therefore interpreted these alternative configurations as long-read sequencing artefacts rather than genuine structural variants. Their occurrence specifically in a repeated region suggests that they are associated with the repetitive structure.

## DISCUSSION

We provide an example of a CNV adaptively introgressed following an anthropogenic marine introduction by characterizing the structural variant previously identified as underlying adaptive introgression in *C. intestinalis*. We first narrowed the 16-gene candidate interval identified by Fraisse et al. (2019) down to a segment containing only three genes, including the candidate gene CYP2U1c. We then showed that the introgressed allele carries a large tandem repeat comprising approximately 23 copies of this segment, with little nucleotide variation among copies. Long-read data further revealed two distinct CYP2U1-containing tandem repeats in *C. robusta*, with different boundaries and copy numbers. Finally, analysis of parent-offspring data confirmed that the CNV follows Mendelian inheritance.

### Benefits and limitations of long-read sequencing for resolving complex tandem repeats

In this study, long-read datasets allowed us to identify the introgressed structural variant as a tandem repeat and to characterize its structure and copy-number variation. Long-read sequencing has enabled the identification and characterization of structural variants (SVs) that remained undetected or poorly resolved with short-read sequencing. For example, adaptive introgression in high-altitude Tibetan populations was initially identified through signatures of selection on single-nucleotide variants (SNVs) linked to the hypoxia-inducible factors EPAS1 and EGLN1 (Huerta-Sánchez *et al*., 2014). The use of long-read data subsequently demonstrated that SVs also contributed significantly to this local adaptation (Quan *et al*., 2021). In line with these recent technological advances, we here revisited an adaptive introgression previously identified with short-read data and used long-read sequencing to characterize the underlying structural variation. While short-read data remain effective for mapping regions targeted by selection, their limitations become apparent when characterizing the structural variants underlying such signals. In our case, using phased short-read data to reconstruct local ancestry across a large sample allowed us to more precisely localize the center of the introgression peak and substantially reduce the candidate interval. However, once the candidate locus was identified as a CNV, a comparison of different coverage-based approaches showed that long-read data provided substantially more reliable copy-number estimates than short-read data. Long-read sequencing also allowed us to examine the internal structure of the tandem array, although its exceptional length remained a major limitation. The shorter repeat alleles identified in a *C. robusta* individual, containing two and five copies, could be fully spanned by individual reads. However, none of our reads fully spanned the high-copy-number introgressed allele. This is not unexpected, as each repeated segment is 5,173 bp long, and an array of 23 copies would extend to over nearly 120 Kb. The long-read datasets of the Roscoff RR *C. intestinalis* individual sequenced by Satou et al. (2021) contained substantially longer reads. By focusing on the longest reads, we identified individual molecules containing up to 23 consecutive copies. However, even these reads did not span the entire array from one unique flanking sequence to the other. Consequently, while we can confirm the presence of at least 23 consecutive copies on a single haplotype, we cannot determine from individual reads if this haplotype contains precisely 23 copies or if the tandem array extends beyond the portion covered by the longest read.

Our study also highlights the challenges of assembling long tandem CNVs comprising many highly similar copies. The NECAT assembly of the Roscoff *C. intestinalis* reference genome further illustrates this difficulty. The 21 copies reconstructed in the assembly differ by no more than one or two nucleotides over 5,173 bp, which provides very little sequence information with which to resolve their order. We therefore examined the underlying reads for features that might help resolve the assembled structure. In particular, copy 14 in the NECAT assembly contains a nested duplication that could potentially provide a unique anchor within the repeat. However, we found no read carrying this duplication at the corresponding position, whether the repeat was explored either from the left or right flanking sequence (Supplementary Fig. S3). We also observed a nested duplication within copy 7 of one read, and a few indels in other reads. Nevertheless, none of these structural variants were found in more than one read. Apart from the nested duplication, copy 14 itself was indistinguishable from other copies in the assembly. We also identified a few reads containing repeated inverted segments (Supplementary Fig. 5). We consider these configurations more likely to represent Nanopore sequencing artefacts than genuine structural variation. The sequence identity among tandem copies may facilitate ectopic interactions between repeated segments during library preparation or sequencing, potentially generating chimeric configurations within individual reads (Spealman *et al*., 2020; Xie *et al*., 2026). While we cannot exclude that NECAT found a valid resolution of the assembly graph and reconstructed a genuine 21-copy haplotype, we were unable to identify information in the underlying reads that would unambiguously support this structure. Thus, while a genuine 21-copy haplotype remains possible, it is not sufficiently supported by the underlying read data.

### Copy number in *Ciona intestinalis*: True variability or methodological biases?

We are confident in our relative copy-number estimates across *C. intestinalis* genotypes (Fig. 2), as well as in our estimate of approximately 23 copies for the introgressed *C. robusta* allele. However, we observed small differences in estimated copy number among individuals. We considered two possible explanations for this variation: genuine biological variation or uncertainty associated with mapping biases and coverage-based copy-number estimation. Genuine biological variation is less likely because it is difficult to explain why the different introgressed haplotypes would have retained such similarly high copy numbers. Given the large number of tandemly repeated units, unequal recombination could occur at many possible positions within the array and, if recurrent, it would generate a broader range of copy numbers rather than consistent differences of only one or two copies (Elemento et al. 2002). Small changes in copy number could potentially arise through other mechanisms, such as replication slippage or transposition. However, we identified no particular sequence motif or repeat feature that would provide an obvious substrate for such mechanisms. Furthermore, such events would not readily explain the striking conservation of the repeat boundaries among haplotypes. We therefore consider estimation uncertainty to be a more likely explanation for the small differences observed among individuals. Copy-number estimates show a slight dependence on average genome-wide sequencing coverage (Supplementary Fig. 1), and the greater variability observed in the short-read dataset illustrates the sensitivity of coverage-based estimates to mapping-related biases (Fig. 2). Although long-read sequencing provides much more reliable relative copy-number estimates across genotypes, estimates derived from read depth remain subject to technical and mapping variation. We therefore interpret the small differences in estimated copy number among introgressed haplotypes as estimation uncertainty rather than evidence for genuine copy-number polymorphism. Consequently, we believe that an ultra-long read approach spanning from one unique flanking sequence to the other would offer the most direct solution for resolving such a long tandem repeat (Jain *et al*., 2018). Furthermore, orthogonal approaches such as droplet digital PCR (ddPCR) could provide independent copy number estimates and have been successfully applied to large tandem repeats (Morton *et al*., 2020).

A single introgressed allele of 23-copy of CYP2U1 is maintained within the *C. intestinalis* species. One hypothesis that could explain why such a large number of copies was favored in the *C. intestinalis* genome is that it is linked to environmental adaptation (Touchard et al. 2024, Le Goff et al. 2026). Indeed, this gene belongs to a large family of genes that encode oxidase enzymes involved, in particular, in cellular detoxification and biotransformation of molecules. Furthermore, a recent study has experimentally demonstrated that CYP2U1 is involved in the inflammatory response in *C. robusta* (Vizzini *et al*., 2021). Thus, pollution could be a key factor in the emergence of the repeat. This hypothesis is also supported by numerous examples of selection driven by human activities that induce rapid adaptation through the development of resistance. This is the case with pesticide exposure, which leads to overexpression of the cytochrome P450 gene in the aphid *Myzus persicae* (Puinean *et al*., 2010), in *Drosophila melanogaster* (Schmidt *et al*., 2010), in *Anopheles funestus* (Wondji *et al*., 2009), and *Spodoptera frugiperda* (Yainna *et al*., 2021). Although all these findings support this hypothesis, a functional study and analysis of expression differences among the various genotypes will be necessary to determine the adaptive role CYP2U1c amplification plays. In particular, the link between tandem repeats and increased gene expression is known in other species (Mileyko *et al*., 2008; Prunier *et al*., 2017), but it needs to be further explored in our case study.

### Variation in *Ciona robusta* and a reasonable evolutionary scenario

The results of Fraïsse et al. (2022) indicate that a high-copy-number CYP2U1c allele is segregating in the *C. robusta* population of Brest Moulin Blanc. Our reanalysis of their short-read dataset showed that three *C. robusta* individuals were consistent with homozygosity for the single-copy allele, whereas other individuals carried a high-copy-number allele similar to the 23-copy allele introgressed into *C. intestinalis*, either at the heterozygous or the homozygous state. Although estimates of absolute copy-number from short reads are imprecise, and copy number in IR heterozygotes was likely underestimated because of differences in mapping efficiency between the divergent genomes, heterozygosity for the high-copy-number allele was independently supported by its segregation in the offspring. Similarly, among the three *C. robusta* individuals analysed with long reads in this study, two were homozygous for a single-copy allele. The persistence of this polymorphism in *C. robusta* suggests that selection on high-copy-number alleles may be local rather than species-wide. In light of the evidence of local adaptation of the introgressed allele in *C. intestinalis* (Touchard et al. 2022; Le Goff et al. 2026), a similar scenario in the donor species could easily explain the coexistence of single- and high-copy-number alleles in *C. robusta*. The selective sweep detected by Fraisse et al. (2022) around this locus in *C. robusta* using SweepFinder (Nielsen et al. 2005) is consistent with a recent intraspecific sweep in the donor species followed by adaptive introgression of the selected allele into the receiving species. Thus, the evolutionary history of this allele may involve two successive episodes of local positive selection, first within *C. robusta* and subsequently, following introgression, within *C. intestinali*s. Furthermore, our long-read data revealed that CYP2U1c variation in *C. robusta* is more complex than the presence or absence of the high-copy-number allele. The third individual carried two distinct expanded alleles, with two copies of a segment containing CYP2U1b, BGLAP and CYP2U1c on one chromosome and five copies of a segment containing BGLAP and CYP2U1c on the other, resulting in a total of seven CYP2U1c and three CYP2U1b copies in the diploid genome. Thus, while the introgressed variation observed in *C. intestinalis* is likely dominated by the single 23-copy allele, *C. robusta* also shows substantial copy-number diversity at this locus. This raises the possibility that selection in *C. robusta* favours increased CYP2U1c copy number, potentially through different segmental duplications, while unequal recombination may have also generated further copy-number diversity over longer evolutionary timescales. In this scenario, the 23-copy-ish allele found at Brest Moulin Blanc would be a local outcome of this process rather than a species-wide optimum. The optimal copy number may also depend strongly on the genetic background. In *C. robusta*, an increased number of CYP2U1c copies may provide a functional advantage, such as an increased gene expression, while a very high number of copies may eventually become costly (Tang & Amon, 2013; Otto *et al*., 2024). Selection could therefore maintain different copy-number haplotypes rather than favouring the largest expansion systematically. Following introgression into *C. intestinalis*, however, this balance may shift. The high divergence between the two species results in strong selection against *C. robusta* ancestry in the *C. intestinalis* genome (Roux et al. 2013; Bouchemousse et al. 2016). A *C. robusta* CYP2U1c allele present at low copy number may therefore provide insufficient benefits to compensate for the cost of introgression into such a divergent genetic background. If increased CYP2U1c dosage is beneficial, a much higher copy number may be required for the benefits to outweigh selection against a suboptimal introgressed sequence The 23-copy allele could therefore have a particularly strong advantage after introgression into *C. intestinalis*, even if such a high copy number is not optimal within *C. robusta*. A shift in the balance between the benefits and costs of increased CYP2U1c dosage across genetic backgrounds could thus explain why diverse copy-number haplotypes persist in *C. robusta*, and why the extreme 23-copy allele was preferentially retained in *C. intestinalis*.

## Acknowledgments

The authors are grateful to the GENSEQ platform of the Research Support Unit MEEB for access to equipment, and to Benjamin Penaud from the Montpellier Bioinformatics Biodiversity platform at ISE-M for his advice for bioinformatic analyses of the Nanopore dataset.

## Author contributions

Manon Le Goff (Investigation, Formal analysis [lead], Writing–original draft [lead], Writing–review & editing [lead]), Fabienne Justy (Methodology, Investigation [lead], Writing–review & editing), Theo Deremarque (Investigation, Writing–review & editing), Frédérique Viard (Conceptualization [lead], Methodology, Funding acquisition [lead], Supervision, Writing–original draft, Writing–review & editing), Nicolas Bierne (Conceptualization [lead], Methodology, Formal analysis [lead], Funding acquisition [lead], Supervision, Writing–original draft, Writing–review & editing).

## Supplementary Material

**FIGURE S1.**
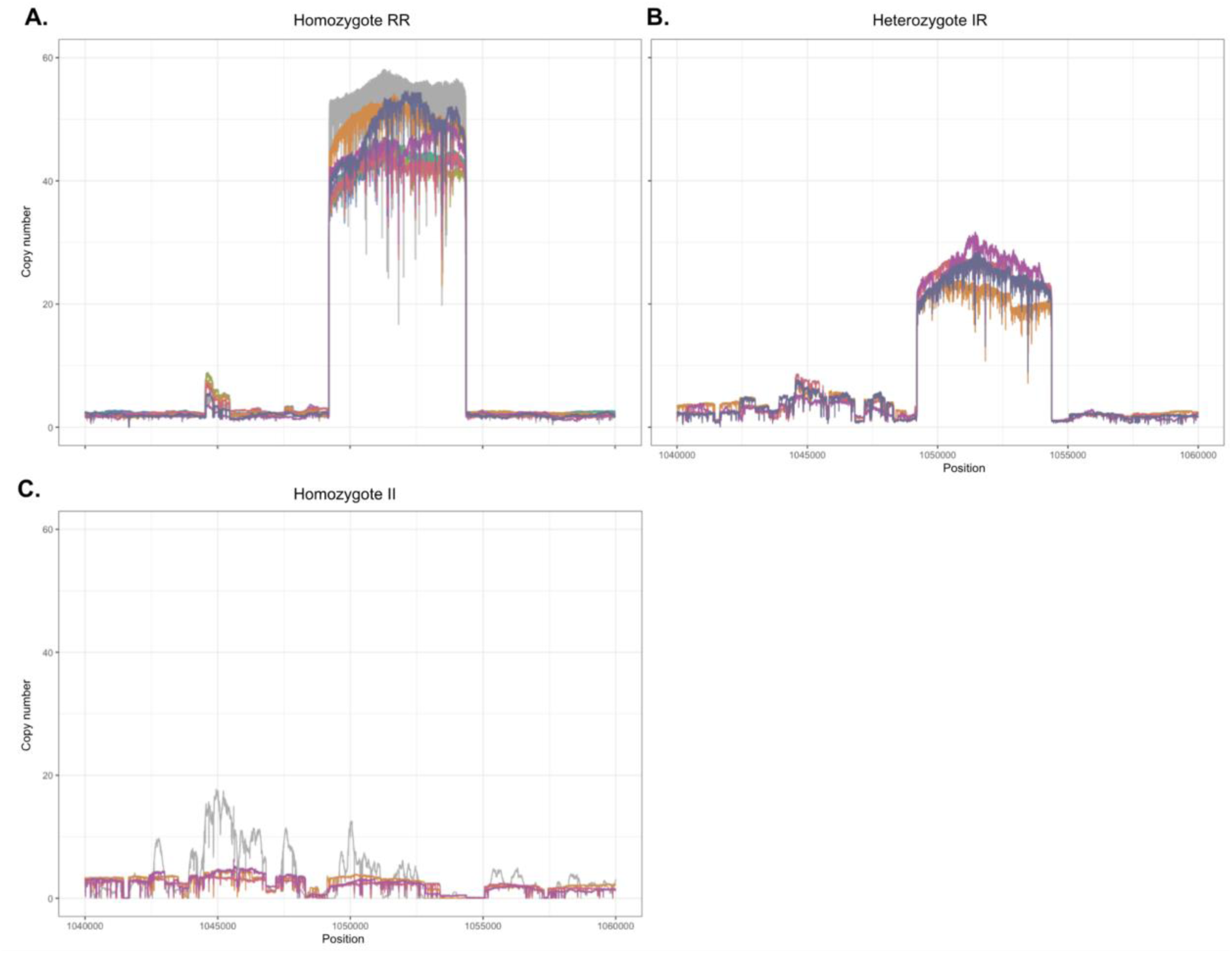
Copy-number estimates based on sequencing coverage for all individuals in the Nanopore dataset. Coverage was normalized by the mean genome-wide coverage of each individual, with a baseline corresponding to two copies. A) Individuals with the RR genotype; the gray curve represents the Roscoff *C. intestinalis* reference genome (Satou et al. 2021). B) Individuals with the IR genotype. C) Individuals with the II genotype; the gray curve represents the Plymouth *C. intestinalis* reference genome (Satou et al. 2021).

**FIGURE S2.**
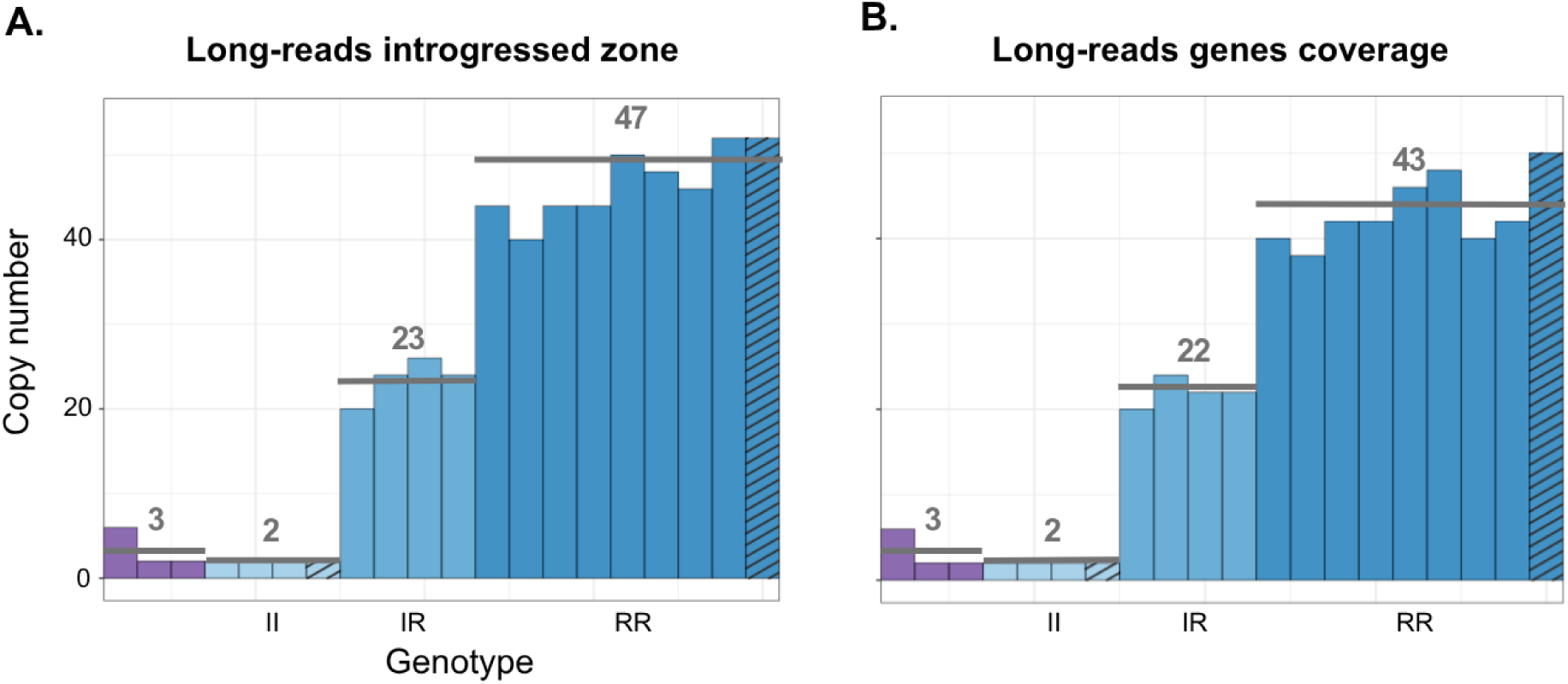
Copy-number estimates according to the denominator used. A) Copy number estimated using the average coverage of the introgressed region (excluding the gene region). B) Copy number estimated using gene-level coverage on chromosome 5 (excluding the gene region).

**FIGURE S3.**
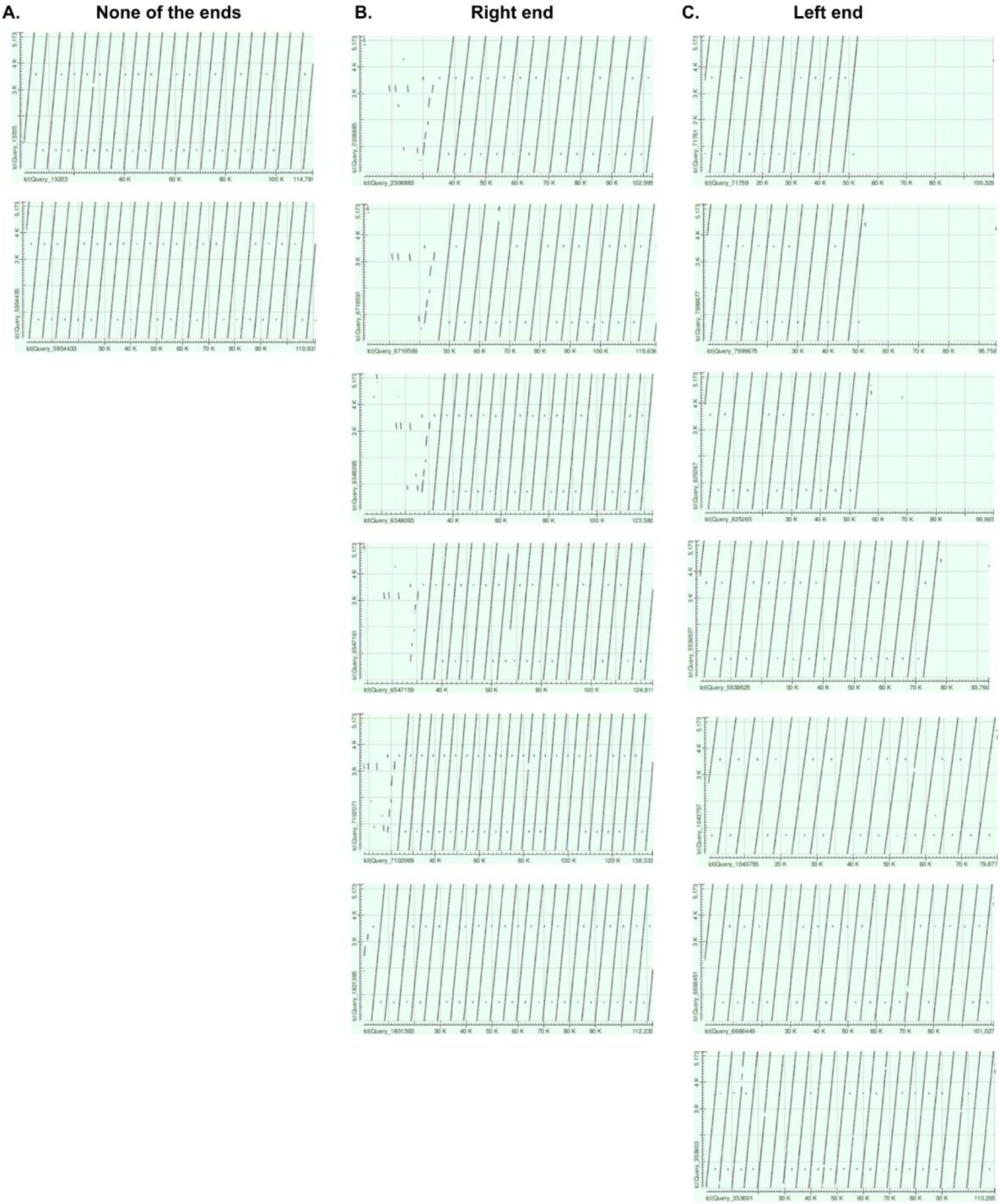
Additional long reads from the Roscoff C. intestinalis dataset (Satou et al. 2021) identified during the search for reads spanning the entire tandem repeat. All dot plots show alignments between individual reads (x-axis) and the 5,173-bp repeated segment (1,049,202–1,054,374 bp; y-axis). A) Reads containing multiple consecutive copies but neither flanking sequence. B) Reads containing multiple copies and the left flanking sequence. C) Reads containing multiple copies and the right flanking sequence.

**FIGURE S4.**
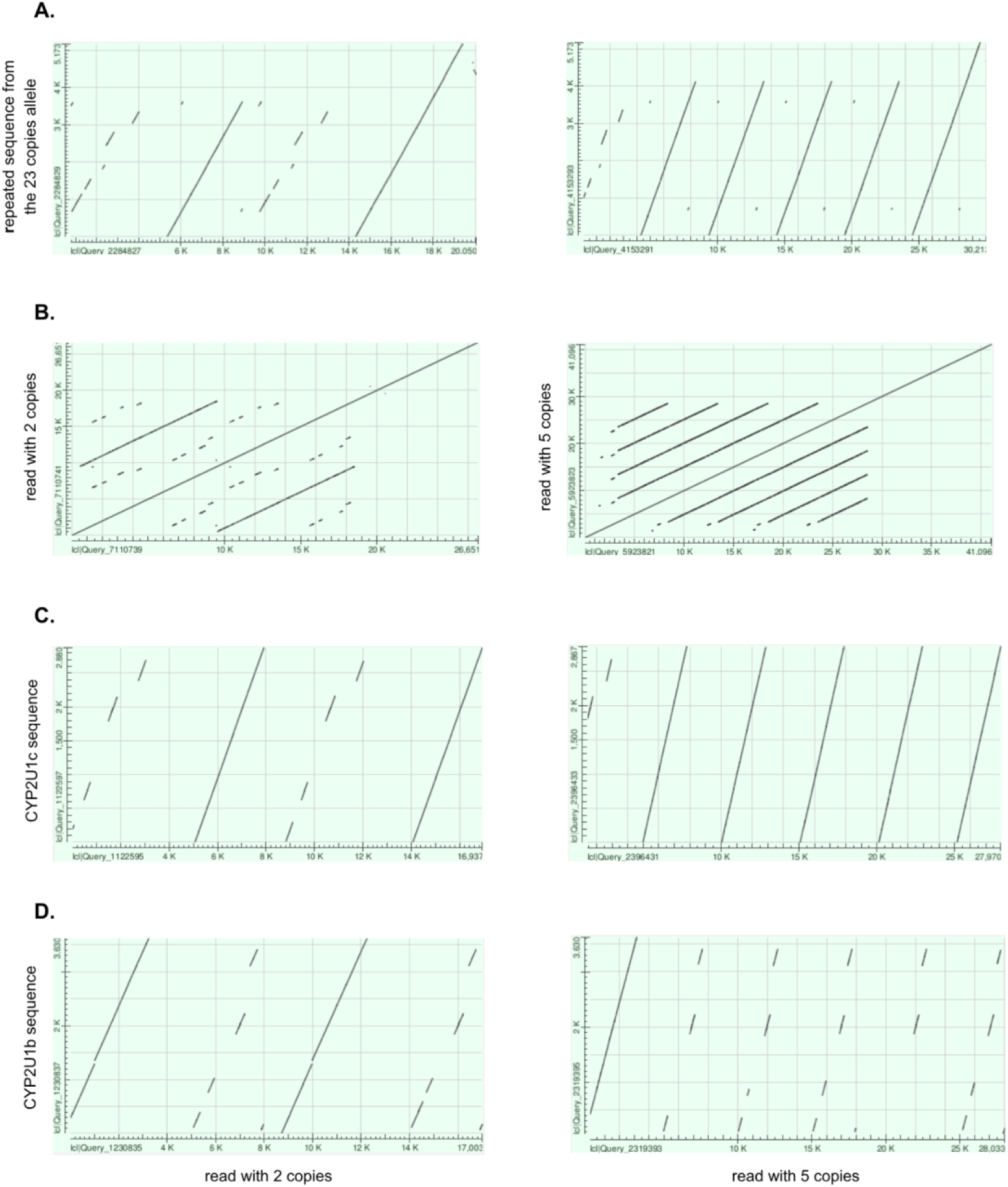
Dot plots of two reads from the *C. robusta* individual Crob_21_ARC_435 used to reconstruct the two haplotypes. In all panels, the left plot shows a read containing a two-copy tandem repeat, whereas the right plot shows a read containing a five-copy tandem repeat. The reads were aligned against different reference sequences to investigate the copy-number profile observed in this individual (Fig. 5). A) Alignment against the 5,173-bp repeated segment identified in *C. intestinalis* (1,049,202–1,054,374 bp). B) Self-alignment of each read, revealing the boundaries of the two- and five-copy tandem repeats. C) Alignment against the CYP2U1c gene sequence. D) Alignment against the CYP2U1b gene sequence. These alignments show that the CYP2U1 genes are repeated in their entirety.

**FIGURE S5.**
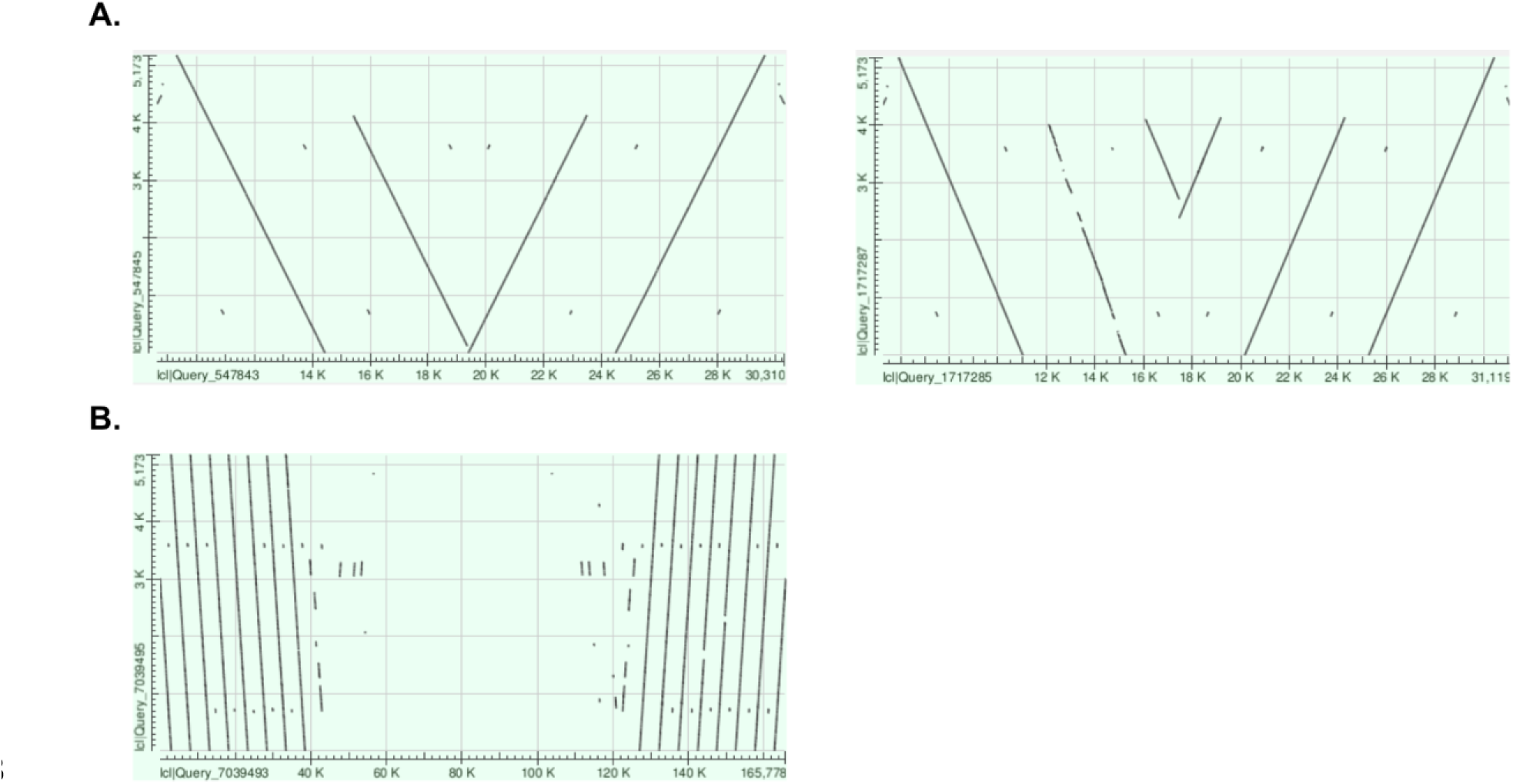
Examples of reads containing inverted duplications. A) Dot plots comparing two reads from the *C. robusta* individual Crob_21_ARC_435 (x-axis) with the 5,173-bp repeated segment (1,049,202–1,054,374 bp; y-axis). Both reads contain two copies followed by two inverted copies, with the second read (right) containing an additional segment. B) Dot plot comparing a read from the Roscoff *C. intestinalis* dataset (Satou et al. 2021; x-axis) with the 5,173-bp repeated segment (y-axis). The read contains 16 copies, comprising eight copies followed by an inverted segment containing eight additional copies.

